# A human cell model of cardiac pathophysiological valvulogenesis

**DOI:** 10.1101/397422

**Authors:** Tui Neri, Emilye Hiriart, Patrick van Vliet, Emilie Faure, Russell A Norris, Batoul Farhat, Julie Lefrancois, Thomas Moore-Morris, Stéphane Zaffran, Randolph S. Faustino, Alexander C Zambon, Yukiko Sugi, Jean-Pierre Desvignes, David Salgado, Robert A. Levine, Jose Luis de la Pompa, André Terzic, Sylvia M. Evans, Roger Markwald, Michel Pucéat

## Abstract

Genetically modified mice have advanced our understanding of valve development and related pathologies. Yet, little is known regarding human valvulogenesis in health and diseases. Genuine human *in vitro* models that reproduce valvular (patho)biology are thus needed. We here developed a human pluripotent stem cell-derived model fit to decode the early steps of human valvulogenesis and to recapitulate valve disease traits in a dish.

Using cellular based, single cell omics-informed and *in vivo*-validated approaches, we derived a population of pre-valvular endocardial cells from a pluripotent stem cell source. These human prevalvular cells (HPVCs) expressed gene patterns conforming to the atrio-ventricular canal (AVC) endocardium signature originally established in E9.0 mouse embryos. In fact, HPVC treated with BMP2, cultured onto mouse AVC cushions, or transplanted into the AVC of embryonic mouse hearts, underwent endothelial-to-mesenchymal transition and expressed markers of valve interstitial cells of different valvular layers demonstrating tissue functionality. HPVCs also differentiated into tendinous/chondrogenic cells in line with the valvular repertoire. Extending this valvulogenic model to patient specific iPS cells, we recapitulated features of mitral valve prolapse and uncovered that dysregulation of the SHH pathway is likely to be at the origin of the disease thus providing a putative therapeutic target.

Human pluripotent stem cells recapitulate early valvulogenesis and provide a powerful model to systematically decipher the origin and lineage contribution of different valvular cell types in humans as well as to study valve diseases in a dish.

## Introduction

Congenital heart diseases are major causes of global mortality in children including Europe and the United States^1,2^. Cardiac valves are affected in up to one third of these life-threatening conditions^3^. Moreover, in adults, the prevalence of valvular diseases dramatically increases with age reaching 13% of the elderly at 75 years of age or older^4,5^. Proper functioning of the valves is indeed essential to ensure efficient blood pumping by the heart. The atrio-ventricular (AV) valves in humans feature two to three leaflets which regulate the direction of blood flow through the mitral and tricuspid sides of the left and right ventricles, respectively. The semilunar valves direct the flow into (pulmonary valve) or out (aortic valve) the chambers. Despite significant progress in the field, there is still limited understanding of disease pathobiology

Mitral valve prolapse (MVP) is among the most common condition and affects 1/40 individuals^6^. Syndromic MVPs are observed in rare and genetically well characterized connective tissues syndromes such as Marfan syndrome, Loeys-Dietz syndrome, or Ehlers-Danlos syndrome. Common non-syndromic MVP is a progressive disease originating from a compromised development of embryonic valves although functional consequences are apparent in middle-aged patients. Genetics studies have identified mutations in selected genes(Filamin A^7^, LMCD1, tensin^8^ and Dachsous^9^ with the disease manifesting by dysregulation of extracellular matrix (ECM) proteins in valve leaflets that precipitates mixomatous degeneration or fibroelastic deficiency. Ultimately valve leakage is the cause of mitral regurgitation^10^. This requires at an advanced stage of the disease surgical valve repair or replacement as no preventive or curative pharmacological alternative is available.

Valve formation is a complex process that begins at HH16 in the chicken, E9.5 in the mouse, and at about 30 days in the human fetus. Intercellular signaling events occurring in the atrioventricular canal (AVC) and the outflow tract (OFT) initiate endocardial-to-mesenchymal transition (endoMT), whereby endocardial cells delaminate and invest the forming cushions. Later in cardiogenesis, this process encompasses interactions between different mesenchymal cell populations, morphogenesis, fusion of cushions, and extracellular matrix (ECM) secretion, leading to the formation of mature and fully functional valves^11^.

The endocardium, as the origin of valvular tissue, is formed by endothelial cells that are distinct from the traditional Flk1^+^ hemoangioblasts^12^. Two paradigms have been proposed to account for segregation of endocardial and myocardial lineages. First, studies in avian embryos^13–15^ suggest that endocardial cells originate in the anterior lateral mesoderm. In line with this concept, *Nkx2.5* deletion in the mouse disrupts endocardial cushions^16^. Second, both endocardial and myocardial cells might share a common multipotent progenitor in the cardiac crescent. *Isl1*^*Cre*^-labeled cells as well as the *Mef2c-(AHF)*^*Cre*^-labeled counterparts give rise to both endocardial and myocardial cells^17,18,19^, thus suggesting that the endocardium also originates at least partially from the second or anterior heart field (AHF) associated with the formation of the right ventricle and OFT, albeit an exclusive AHF origin has been challenged. In fact, a contribution of endothelial cells derived from MesP1+ cells to the formation of the endocardium was proposed^19^.

Genetically modified mouse models have provided important clues as to the cell lineages and signaling pathways that contribute to valve formation^20–23^. However, these murine transgenic models are limited in their potential to reveal mechanisms underlying complex processes, as aspects of signaling interaction, cell metabolism, epigenetics, and mechano-transduction are difficult to mechanistically separate *in vivo* and might not be identical in human valve development. In fact, cell lineages that contribute to the valves in humans have been elusive and have only been inferred from knockout mouse models recapitulating part of the human pathophenotype. A human specific cell model of valvulogenesis that could be extended to cells derived from individual patients would definitively advance the understanding of developmental mechanisms driving valvulogenesis in health and disease.

Pluripotent stem cells have been reported to recapitulate early developmental processes including cardiac myogenesis^24^ but differentiation of these pluripotent cells towards valvular specific cells has not yet been reported. Here, leveraging embryology data across species, we report that human pluripotent stem cells are able to recapitulate the developmental process of valvulogenesis. We used a population of human pluripotent stem cell-derived MESP1^+^ sorted cardiovascular progenitors^25^ and directed their fate concomitantly towards the first/second heart field and endothelial cell lineages. We report that these cells collectively or at the single cell level express a set of salient genes that mark pre-EMT (E9.0) mouse AVC. These cells undergo EMT in both notch-dependent and independent manners and express specific valve proteins when treated with BMP2, or when seeded onto collagen hydrogels, or grafted *in vivo* in mouse embryos. Furthermore using patient specific induced pluripotent stem (iPS) cells harboring a mutation in Dachsous^6^ and in turn propensity to mitral valve prolapse^9^, we recapitulated cell features of the valvulopathy, and uncovered the molecular mechanisms of the disease that points to a therapeutic target. Human pluripotent stem cell-derived endocardial cells thus represent a novel model of human valvulogenesis enabling future studies on mechanisms of human valve pathogenesis.

## Materials and Methods

### Cell culture and sorting

HUES cell lines HUES-24 and HUES-9, obtained from Harvard Stem Cell Center (Dr Chad Cowan), were previously described^25^. HUES-24 was here used to generate a transgenic cell line expressing GFP under the control of the *Sox9* promoter. To engineer the DNA construct, the 298-bp minimal *Sox9* promoter and two cis-regulatory elements^26^ were excised from the pGBW101-1,2 vector (kindly given by Dr Bien-Willner, Baylor College of Medicine, Houston) and subcloned in a pAcEGFP1-1 vector. The DNA construct was electroporated and cells were selected with G418 for 10 days. HUES cell lines were cultured on Mouse Embryonic Fibroblasts (MEF) prepared from E14 Mouse embryos using KO-DMEM medium supplemented with β-mercaptoethanol, glutamine, non-essential amino acids, 15% KOSR and 10 ng/ml FGF2.

To generate a MesP1^+^ cell population^25^, human pluripotent stem cells (both HUES and iPS cells) cultured on MEF were treated for 4 days with wnt3a (100 ng/ml) and 10 ng/ml BMP2 in the presence of 1 μM SU5402, a FGF receptor inhibitor, in RPMI/B27. For sorting, trypsinized cells were incubated for 30 min with gentle occasional agitation with EasySep™ Human Whole Blood CD15 (SSEA-1) Positive Selection Kit (Stem cells technology) (25 μl/ml suspension cells) in D-PBS supplemented with 0.5% (wt/vol) Bovine Serum Albumin (BSA) and 2 mM EDTA at room temperature. Cells were then transferred to a magnet. Cells were washed three times with 5 ml D-PBS-BSA/EDTA. Immunofluorescence, using an anti-CD15-FITC (Amicon) antibody, carried out directly after sorting revealed 90% purity of SSEA-1^+^ cells (data not shown). HUES or iPS cells derived SSEA1+ sorted cells were phenotyped by RT-real time PCR and expressed *MESP1*, *MEF2C*, *NKX2.5*, *ISL1* but not *CD31*, *ENG* or *CDH5*. Immunostaining documented that 90% of cells were MESP1^+25^.

To direct differentiation of (CD15+)SSEA-1^+^ cells towards endocardial HPVC cell fate, cells were cultured on MEF plated at low density (10000 cells/cm^2^) on fibronectin-coated plates and treated with 30 ng/ml VEGF, 10 ng/ml FGF8 and 2 ng/ml FGF2 for 6 days and sorted with anti-CD31 conjugated magnetic beads (Miltenyi, France). Separately, CD31+ cells cultured on fibronectin-coated plates were further induced to undergo EMT using 100 ng/ml BMP2 for 2 days in the presence or absence of DAPT (1 μM). In a series of experiments, cells were transfected with a NCDI (Notch intracellular domain) expression plasmid (1 μg) or empty backbone vector using lipofectamine 2000.

### iPS cells

iPS cells were derived using the Sendai viral vectors (kit Lifetech themofisher France). Cells were characterized by immunostaining OCT4 (anti-OCT4 santa-cruz sc-9901), Sox2 (anti-SOX2 santa-cruz) and NANOG (anti-NANOG R&D). Cells were then differentiated toward endoderm by treatment with 100 ng activin in DMEM supplemented with 10% FCS for 3 days, mesoderm by adding CHIR-99021 (5μM) the first day, CHIR-99021 and BMP2 (10 ng/ml) the second day and IWR1+ BMP2 the third day. Ectodermal cells were obtained by culturing cells for three days in RPMI + N2 supplement and 0.5 μM retinoic acid. Genomic integrity was tested at passages 15-20 using digital PCR of copy number variants of main human recurrent genomic abnormalities (stemgenomics, Montpellier, France).

### Reverse-Transcription Quantitative Real-Time PCR (Q-RT-PCR)

RNA was extracted from SSEA1^+^ cells or VEGF/FGF8-treated using a Zymo research kit. RNA (1μg) was reverse-transcribed using the Superscript II reverse transcriptase (Invitrogen, Cergy, France) and oligo(16)dT primers. Q-PCR was performed using SYBR Green and a Light Cycler LC 1.5 (Roche Diagnostic). Amplification was carried out as recommended by the manufacturer. Data analysis and primers specific for human genes are described previously^25^ and in supplementary table 1.

### Microarrays

AVC and the primary ventricle of E9.0 mouse embryos were dissected out, and RNA (triplicates) was extracted with a Zymo research kit. RNA was also extracted from HPVCs. cRNAs were profiled using Illumina Mouse WG-6 v2 BeadChips or human Expression HG-U133 arrays and analyzed in Genespring GX 11.0. Data were quality filtered to exclude signal intensities below background, and expression profile differences of 4 fold or greater at a significance threshold of 0.01 or less were compiled into upregulated and downregulated transcript lists. Bioinformatically filtered genes were used for downstream analysis in Ingenuity Pathway software to examine network relationships as well as identify overrepresented gene ontologies.

Cluster analysis of AVC and mesenchymal stem cell datasets were conducted as follows. Previously published AVC (GDS3663) and mesenchymal array datasets (GDS1288) were downloaded from GEO. HPVCs gene expression profiles were examined using HG-U133 arrays and signal values were computed using GCRMA normalization. All expression signals for each dataset were Log_10_ transformed and normalized by z-score transformation^27^. Experimental replicates were then averaged for each study, human and mouse orthologs and/or homologs between microarray data sets were determined by matching NCBI gene symbols across experiments and between species. This resulted in 9,385 orthologues that were then clustered using the HOPACH algorithm for gene clustering with the cosangle distance metric^28^ or by hierarchical clustering with the Euclidean distance metric.

### Single cell-sequencing

iPS cell-derived HPVC and and iPS cell-derived valvular interstitial cells were dissociated with trypsin in single cells and processed in Chromium (10X genomics). cDNA libraries were sequenced with a Next-seq Illumina sequencer. A first analysis was performed with cell Ranger and C-loop 10X genomics softwares. Clusters and subclusters were defined using genes differentially expressed in cell clusters in comparison with all other cells with a threshold of Log_2_ equal to at least 2. Then a secondary analysis was done using the SCDE package^29^. 1895 genes were detected as expressed in 50% of cells and 4576 in 25% of cells (Supplementary Fig 1).

**Figure 1:**
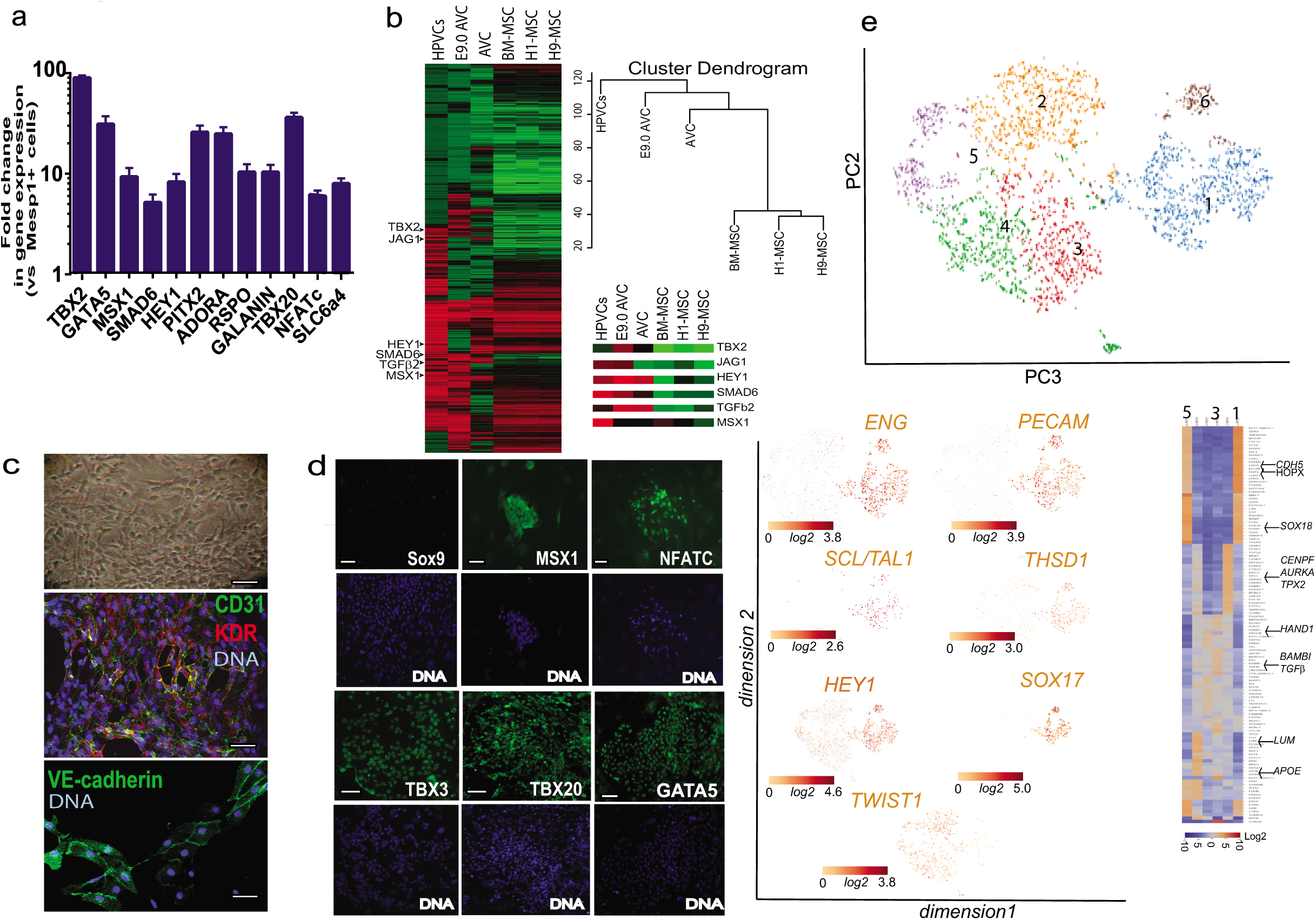
Gene and protein expression profiles of human valve progenitors. SSEA-1^+^-sorted human MESP1^+^ cells were treated with 10 ng/ml FGF8 and 30 ng/ml VEGF for 6 days. (a) RNA was then collected and cDNAs of FGF8/VEGF treated cells (HPVCs) were run in Real Time PCR (mean±SEM of 9 experiments). Data are normalized to 1 as the level of gene expression in SSEA1+MESP1+ cells. (b) cRNAs (n=3 experiments) were used for microarrays and normalized *versus* MESP1^+^ cells from the same respective cell sorting. Heatmaps of transcriptomes of HPVCs, E9AVC (our data), AVC GDS3663 and MSCs (GDS1288). A few AVC specific genes are highlighted in the inset. (c) Bright field image (top) and co-immunostaining of VEGF/FGF8-treated SSEA-1^+^/MESP1^+^ derived colonies with anti-CD31 and anti-Flk1(KDR) or anti-VE-cadherin antibodies. Data are representative of 5 experiments. (d) Immunostaining of VEGF/FGF8-treated SSEA-1^+^/MESP1^+^ derived colonies with anti-Sox9, -Msx1, -Nfatc1,-Tbx3, -Tbx20 and -GATA5 antibodies (green) and DAPI (blue). Data are representative of 5 experiments. The scale bar indicates 50 mm. (d)(e) HPVCs were further sorted using anti-CD31 conjugated beads and used in single-cellRNA sequencing. t-distributed stochastic neighbor embedding (t-SNE) 2D cell map 10X genomics (n=2440 cells) (upper panel). Highlight of cell populations expressing genes marking endothelial, hemogenic and early EMT cells (lower panel) and heatmap of graph-based Log2 fold changes in gene expression of cell cluster compared to all other cells (lower right panel)

### Antibodies

Antibodies used for cell or embryo immunofluorescence were raised against Isl1 (Developmental Hybridoma bank, Iowa University), Tbx20 (novus biological H00057057-B01), Msx1 (Abcam ab49153-100), Tbx3 (Santa Cruz sc-17872), aggrecan (Chemicon AB1031), versican (Chemicon AB1033), smooth muscle actin (Sigma A-2547), Filamin A (Epitomics, CA, USA), vimentin dylight 550 (Novus biological), periostin (Abcam ab140141), CD31 (BD pharmigen), VE-cadherin (R&D systems/MAB9381), Sox9 (a gift from Pr Wegner, University of Nurnberg, and Santa-Cruz Sc17431), GATA5 (Abcam ab11877), Hyaluronan binding protein (Millipore 385911), collagen I (Abcam 34710), anti-Patched1 (Merck-Millipore, France), human Lamin A/C (Novacastra), and anti-NFATc (santa-cruz H110). Immunostaining of cells and paraffin-embedded embryonic heart sections was carried out as previously described^25^.

### Cell imaging

High content imaging was performed in 96 wells plates using an Arrayscan (Cellomics Thermo Fisher Scientific) attached to an inverted microscope (Carl Zeiss, Inc.) using 20x N-Achroplan objective, NA 0.45, at room temperature. Other images were observed in epifluorescence microscopy (Zeiss microscope) or using a Ultraview Vox Spinning disk Perkin Elmer confocal microscope driven by the Volocity software or using a TRIO multispectral analysis setup (Caliper, Science). To visualize cilia staining, stacks of images were acquired using a Zeiss observer epifluorescence microscope. Then the images were deconvoluted using Autoquant and reconstructed in 3D using Imaris software (IMARIS). Quantification of extracellular matrix proteins was performed by thresholding images acquired with a Zeiss observer microscope equipped with a Colibri light source using the same LED intensity and same exposure time for the CCD camera for all images. The surface of the cell field labelled by the antibody was calculated using Image J (NIH image).

### AVC and OFT explants

Dissection and culture of explants on collagen I hydrogel were carried out as described^30^. Briefly, after dissection, the OFT or AVC explants were cultured with the endocardium facing the gel in DMEM/ITS medium.

### Tendinous/chondrogenic differentiations of HPVCs

Chondrogenic differentiation was performed according to established methods^31^. HPVCs were isolated from MEF using collagenase IV (Life Technologies, France), transferred into 15 mL of polypropylene centrifuge tubes (500,000 cells/tube) and gently centrifuged. The resulting pellets were statically cultured in DMEM high-glucose medium with glutamine, penicillin/streptomycin and chondrogenic supplements (1X insulintransferrin-selenium, 1μmol/L dexamethasone, 100 μmol/L ascorbic acid-2-phosphate), and 10 ng/mL TGF-β1. After 3 weeks, RNA was extracted from pellets and subjected to Q-RT-PCR. Cell pellets were fixed in 4% paraformaldehyde overnight, embedded in paraffin, and sectioned in 5 μm slices. Sections were stained with anti-collagen1a and –Sox9 antibodies.

### Collagen gel culture

SSEA-1^-^ HUES cells, SSEA-1^+^ HUES cells or SSEA1^+^ HUES cells treated with VEGF and FGF8 and FGF2 (HPVCs) were cultured alone and aggregated. Preparation of collagen gel (1 mg/ml type I collagen from rat tail tendon, BD Sciences) was described^32^. Resultant cell aggregates were placed on hydrated collagen gels and cultured for 48-72 h.

### Cell injection in embryos, embryo culture, and isolated embryonic heart culture

Mouse embryos were collected from E10.5 pregnant mice and cultured in DMEM or in M2 medium supplemented with 10% FCS at 37°C and set with pins (isolated heart culture) or between pins (whole embryo culture) in a culture dish filled with silicone (Figure 4A). Glass injection pipettes were pulled with a P-87 Flaming/Brown Micropipette Sutter Puller to get a 20 μm tip inner diameter. To optimize injection, a pipette was filled with the green dye CDCFDA, SE (Invitrogen/Molecular Probes, USA) and the dye injected into AVC.

Micropipettes were filled with HUESC-derived HVPC, or MesP1^+^ SSEA1^+^ or SSEA1^-^ cells as controls in DMEM-10% FCS medium at a concentration of 10^5^ cells/μl. One to three injections at a pressure of 200 hPa for 1 s were carried out in both the wall and the lumen of the AVC or OFT. Cell injections in brain were used as a negative control. At 4 h following injection, hearts were dissected and cultured at the air-liquid interface on insert coated with matrigel and set in multiwells plates filled with DMEM-10% FCS. After 48 h culture, embryonic hearts were PFA-fixed and embedded in 1% agarose blocks prior to paraffin embedding. Alternatively, cells were injected in the AVC of E9.5 embryos through the yolk sac and embryos cultured in rolling tubes in 25% M16 medium /75% rat inactivated serum and gassed with 40% O_2_/5%CO_2_ for 24-48 h^33^.

### Statistics

Data are expressed as mean±SEM. Experiments were repeated up to 9 times. Student-t-test was used to compare datasets after checking for continuous probability distribution (Gaussian distribution) for each data group.

## Results

### Human pluripotent stem cells differentiate into endocardial prevalvular cells

We developed here an original protocol to differentiate human ES or induced pluripotent stem (iPS) cells into genuine valvular cells. Undifferentiated pluripotent stem cells were first differentiated into MesP1^+^ cardiovascular progenitors using Wnt3a (100 ng/ml) and BMP2 (10 ng/ml) for two days and then BMP2 alone; cells were then sorted using the BMP2-induced SSEA-1 cell membrane antigen^25^ and plated on MEF in fibronectin-coated plates. To segregate myocardial and endocardial cell lineages from the MesP1^+^ cell population, cells were treated with VEGF (30 ng/ml), an inducer of endothelial cell fate. We further added FGF2 (2 ng/ml) and FGF8 (10 ng/ml) for 6 days. FGF8 was used to trigger mesodermal cell fate toward endocardial cells at the expense of myocardial cells^34^.

Cells were then phenotyped using real time PCR, immunofluorescence and high content imaging. Figure 1a shows that VEGF/FGF8 treated cells intensely expressed *TBX2, TBX20, GATA5, MSX1, SMAD6, PITX2, HES1, NFATc, ADORA* and *GALANIN,* when compared to the level of expression of these genes in the MESP1^+^-sorted cell population. A global transcriptomic signature of the human endocardial cells using gene arrays further revealed enrichment of VEGF/FGF8 treated cells (or Human PreValvular Cells, HPVCs) *vs* MESP1^+^ cells in *TGFβ1* and *TGFβ2, MSX1*, *THROMBOSPONDIN*, *PITX2, ERBB4, ADAM19*, and *CAV1,* expressed in the mouse AVC endocardium^35,36^, *PDGFRα*, *LMCD1* and its target *GATA6*, enriched in cardiac cushion at the level of mRNA^37^ as well as in endothelial genes *KDR*, *FLT1*, *FLI1, TIE2, NFATc* (Fig. 1b; supplementary figure 2, transcriptomic data GEO dataset). Conversely, *MESP1*, and cardiac chamber specific genes *TBX5*, *MEF2c*, *NPPA, MLC2v* were not expressed in human prevalvular cells (HPVCs) (data not shown, see transcriptomes). Comparisons of HPVC transcriptomes with H1 and H9 ESC lines-derived mesenchymal cells or bone marrow derived mesenchymal cells as well as with E9.0 AVC cells indicated HPVC clustered with E9.0 AVCs and to a lesser extent to previously reported AVCs^38^ but displayed little correlation with human ESC-derived or mesenchymal stem cells (Fig.1b). The HPVC transcriptome further showed the presence of genes specific to AVC (*TBX2*) and endocardium (*MSX1*), involved in Notch signaling (*JAG1, HEY1)* or in endocardial cushion mesenchymal transition (*TGFβ, SMAD6*) (Fig.1b).

**Figure 2:**
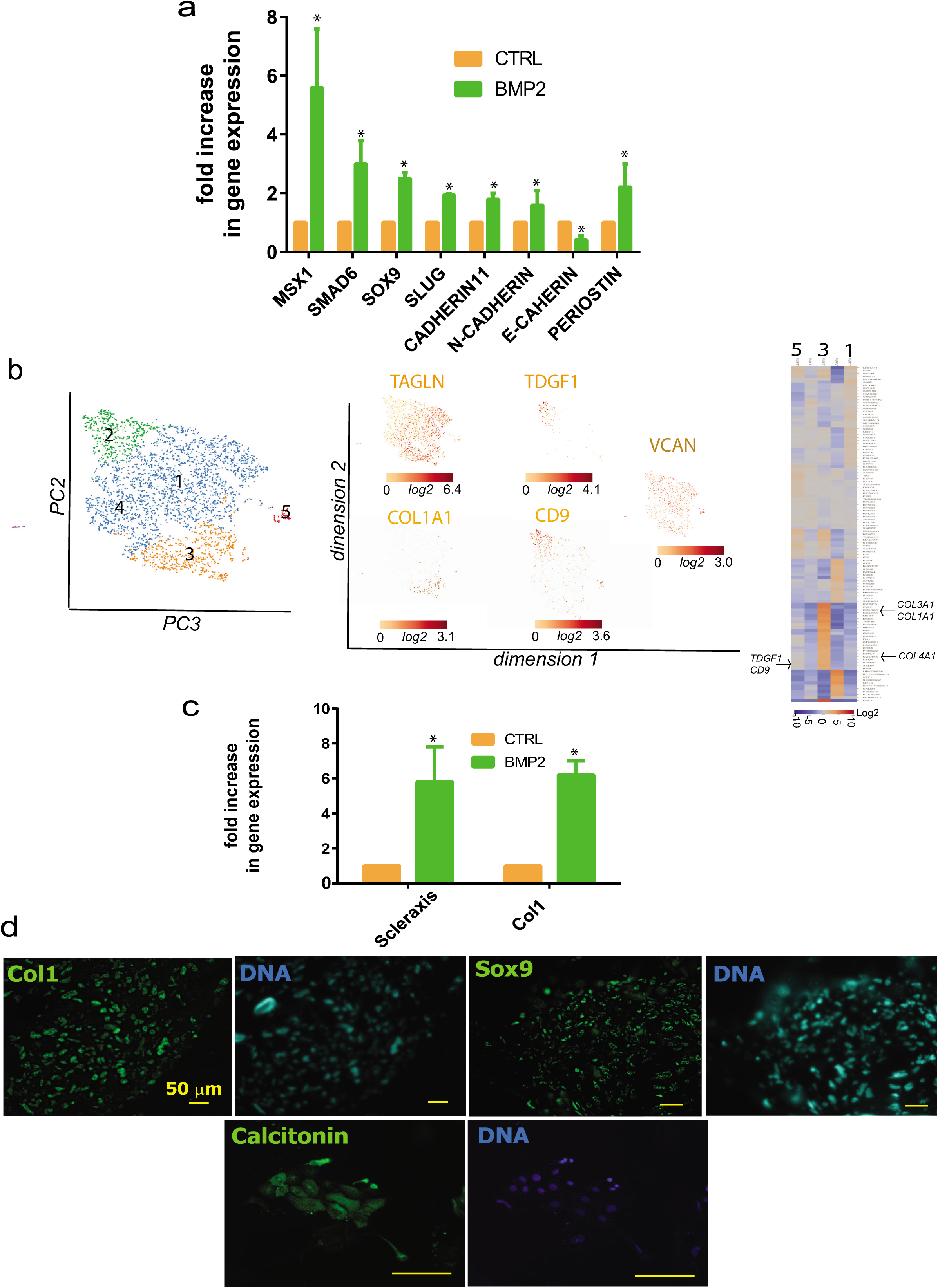
EMT of HPVC cells. (a) After 1 week of FGF8/VEGF treatment on MEFs, valve progenitors (HPVCs) were recovered with trypsin, seeded on fibronectin-coated wells and treated with 100 ng/ml BMP2. After 2 days, RNA was recovered and cDNAs were run in real-time PCR for post EMT markers. BMP2 samples are normalized on control (before treatment) samples, showing an increase in the expression of post-EMT markers (*) (b) t-distributed stochastic neighbor embedding (t-SNE) 2D cell map 10X genomics (n=3700) cells (left panel). Highlight of cell populations expressing *TAGLN* and genes marking more specifically fibrosa (*COL1A1*) or spongiosa *(CD9, TDGF1*) (middle panel) and heatmap of graph-based Log2 fold changes in gene expression of cell cluster compared to all other cells (right panel). (c) **HPVCs give rise to tendinous/chondrogenic cells**. HPVCs were pelleted into a tube and treated with a chondrogenic medium as described in the methods. Real Time PCR shows an up-regulation of Scleraxis and Collagen 1a (Col 1) genes versus non-treated cells following three weeks of treatment (two experiments in duplicate; (*) significantly different p≤0.01). The cell pellets were embedded in paraffin and 5 um sections stained with anti Sox9, anti-Collagen1a and anti-calcitonin antibodies.Data are representative of 3 experiments. The scale bar indicates 50 mm.

HPVCs featured a morphology of endothelial cells. Immunofluorescence and high content imaging confirmed that expression of endothelial CD31 and KDR was turned on in 95% of cells (out of 850 scored cells in 2 separate experiments), VE-cadherin (Fig. 1c), as well as AVC and endocardial cells (GATA5, TBX3, TBX20, NFATc and MSX1) protein markers (Fig. 1d).

To estimate the cell heterogeneity of HPVCs, a single cell-sequencing approach was used. CD15+ Mesp1+ sorted cells were plated on fibronectin coated dishes and induced with VEGF/FGF8/FGF2 to an endocardial cell phenotype. Cells were then further sorted using an anti-CD31 antibody and used for single cell-sequencing. We performed a two-step data analysis to cluster cells by principal component analysis. Figure 1e revealed the endocardial phenotype of 3000 HPVCs at a single cell level. Although the cells were CD31-sorted, the cell population was heterogeneous. Five main cell clusters were identified as expressing a specific gene pattern depending upon their stage in the early process of EMT (Fig. 1e).

One third of cells (1105) (cluster1) were *CDH5*+ including endoglin (*ENG)+* cells (849) and *PECAM1+* cells (729) as well as *NOTCH4*+ (1005) cells, pointing to an endothelial and endocardial cell population. These cells also highly expressed *SOX17, SOX18*, *KDR* and *ETS* indicative of the endocardial phenotype (supplementary table 2). *TWIST1*+ cells clustered as a mirror of ENG+ cells although some cells were *ENG+TWIST1+.* 90% of *TWIST1*+ cells were negative for *SNAI1* but still positive for *ETS1+,* an endothelial gene confirming the early EMT stage of these cells. *THSD1* as well as *HEY1* AVC endocardial genes were both enriched in the *CDH5+* cell population (Fig.1e) while *GATA5, TBX2, TBX3*, *MSX1* were found expressed in both endocardial and *TWIST1+* cells (supplementary table 2).

Interestingly, a cluster of cells (180 cells within cluster1) expressed high level of *ETS1, EGFL7, HOPX* and *SCL/TAL1* suggesting the presence of a hemogenic endocardial cell population^39^ (Fig. 1e). Cluster 2 included highly proliferative cells expressing genes involved in cell mitosis such as *GTSE1, CENPF, AURKA, BIRC5* and *TPX2*. Cluster 3 included cells expressing genes of the TGFβ signaling pathway (*BAMBI, TGFβ1)* and *PDGFRα.* Cluster 5 included cells more advanced in the EMT process expressing among others *APOE, high level of LUM, POSTN, and ACTA2*.

Thus, human ESC derived HPVCs faithfully reflected the phenotype of endocardial cells before and at the onset of EMT within the AVC and showed robust correlation of gene expression with E9.0 AVC cells rather than with any other mesenchymal stem cell type.

### Human valvular progenitors undergo EMT and depend on Notch signaling in vitro

To test the potential of pluripotent stem cell-derived HPVCs to undergo EMT, a key process executed by endocardial cells to form cardiac cushions *in vivo*, HPVCs were further treated with BMP2 (200 ng/ml) for 48 h. Gene expression was monitored by real time PCR. When compared to non-stimulated cells, the level of expression of *MSX1*, *SMAD6*, *SOX9*, *SLUG*, *CADHERIN 11*, *N-CADHERIN* AND *PERIOSTIN* was significantly increased while *E-CADHERIN* was decreased. (Fig. 2a).

Single cell-sequencing analysis was further performed after BMP2 treatment. Clustering within 5 tight groups revealed that most cells (2598) expressed *TGLN,* a myofibroblast marker enriched in the ventricularis layer of the valve^40^. Cells initiated specification into fibrosa, spongiosa or ventricularis layers of the valve. The *COL1A1* cell cluster (cluster 3) was enriched in most collagen genes (*COL1A1, COL1A2, COL4A1, COL6A2, COL3A1)* as well as in genes enriched in fibrosa such as *BGN*^*40*^. Cluster 2 was enriched in genes *TDGF1, CD9, VCAN, and ALDH2* found in the spongiosa^40^. Cluster 1 included endothelial cells still expressing CDH5 and PECAM1 (Fig. 2b; supplementary table 3).

Notch has a crucial function in the process of EMT in cardiac cushions^41–43^. We thus tested the role of the Notch pathway in BMP2-induced EMT of HPVCs. BMP2-induced expression of *SLUG* and *PERIOSTIN* was inhibited by 1 μM DAPT (N-[N-(3, 5-Difluorophenacetyl)-L-alanyl]-S-phenylglycine t-butyl ester) (Supplementary Fig. 3a), a γ-secretase inhibitor that blocks Notch pathway activation, indicating that expression of these two markers is Notch-dependent. Activation of the Notch pathway following transfection of Notch intracellular domain NICD strongly turned on the expression of *PERIOSTIN* as well as of *SLUG* and *MSX1* (Supplementary Fig. 3b), suggesting that as reported *in vivo*^41, 42^ Notch regulates EMT via SNAIL and SLUG activation.

**Figure 3:**
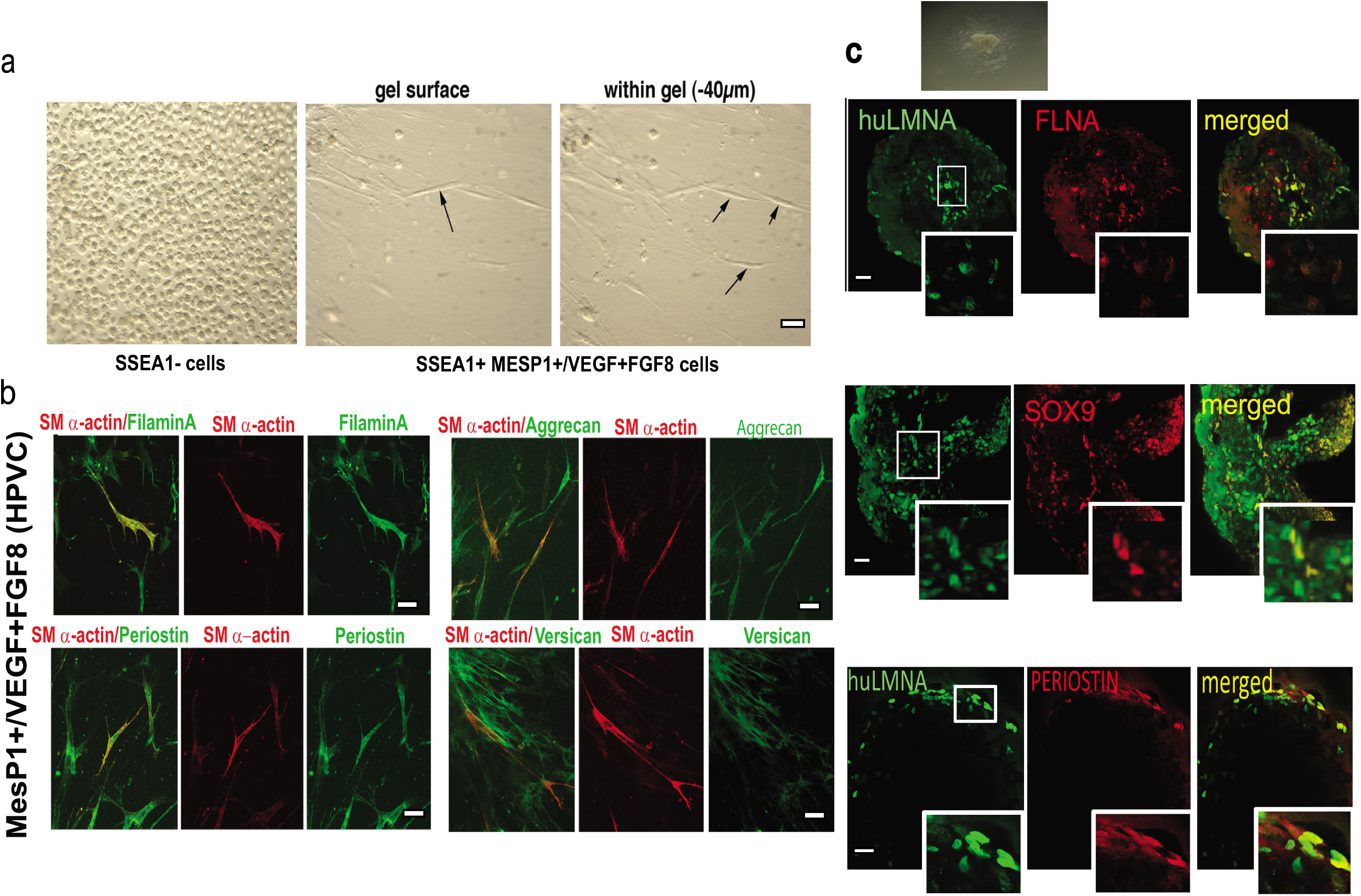
Ex vivo differentiation of HPVCs. (a): <u>Left panel</u>: After 4 days of BMP2 treatment MESP1^-^(SSEA1^-^) cells were collected and cultured in hanging drops. Cells were able to form aggregates, but when plated on collagen I hydrogels they did not adhere or invade the gel. <u>Middle panel</u> After SSEA1^+^ sorting and treatment for 1 week on MEFs with FGF8 and VEGF, 40,000 valve progenitors were cultured as hanging drops for 18 h and plated as aggregates on collagen I hydrogels. In 48 h, invasive mesenchymal cells were seen in the gel (−40 μm from the surface), fixed and immunostained for post-EMT markers SM alpha actin together with filamin A, periostin, versican, and aggrecan. (b): 20,000 MESP1^+^, or MESP1^-^ progenitors or FGF8/VEGF-treated MESP1^+^ progenitor cells were mixed with 20,000 freshly isolated chick H-H stage-24 AV cushion mesenchymal cells to aggregate. Cells were cultured with or without BMP2 for 48 h and stained for human LaminA/Cantibody (green). Very few SSEA1^+^ and SSEA1+^-^ cells adhered and survived whereas FGF8/VEGF treated cells compacted themselves at the middle of the chicken-human cell aggregate and, when triggered with BMP2, some cells started to spread out from the aggregate. (c): FGF8/VEGF treated MESP1+ progenitors were placed on the top of E9.5 mouse AVC explants on collagen gels (inset), cultured for 2 days and co-immunostained with anti-human LaminA/C and anti-Sox9, -filamin A or -periostin antibodies. Bar scale indicates 50 μm. Data are representative of at least 6 separate experiments.

**Figure 4:**
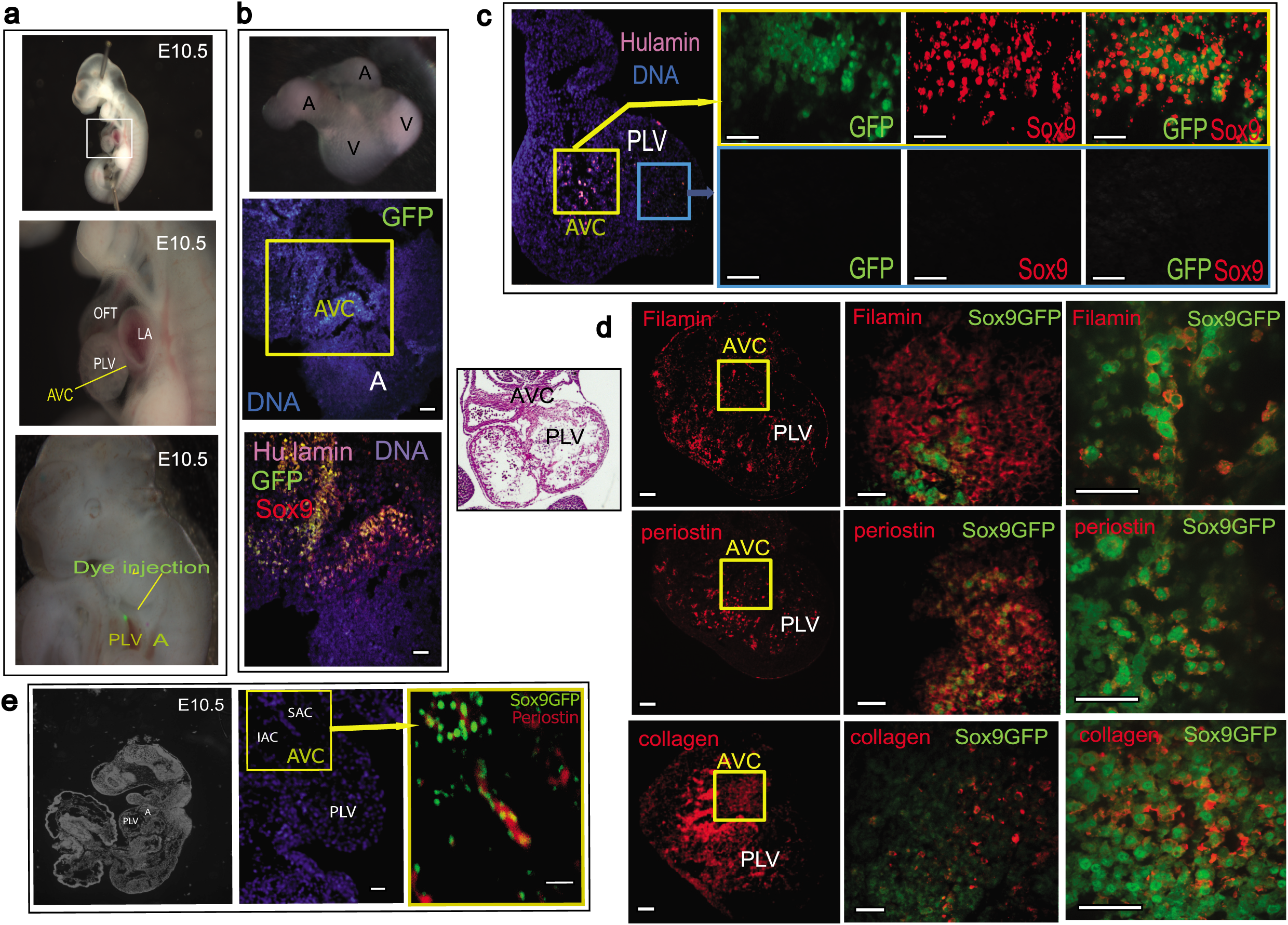
In vivo differentiation of HPVCs in mouse embryonic heart. SSEA1^+^ MESP1^+^ *Sox9*GFP cells were sorted and treated with VEGF/FGF8 for 7 days before injection into E10.5 mouse AVC. (a) An E10.5 embryo is stretched to expose the AVC. Dye injection shows the target of the injection. (b) Two hours after injection the hearts were dissected and placed in culture for two days on matrigel coated inserts (top inset). The heart sections were immunostained with anti-human LmnA/C, –GFP, and –Sox9 antibodies. (c) Two sites of injection and resulting cell fate. Cells were injected into the AVC or the chamber. Only the cells injected into the AVC (visualized on the eosin/hematoxylin stained heart section in the let inset) underwent EMT and expressed Sox9-GFP. (d, e) Sox9GFP VEGF/FGF8 treated cells were injected into the AVC and the hearts were cultured for two days (d) or the whole embryos were cultured in their yolk sac for 36 hours. The heart sections were stained with anti-filaminA, -periostin or –collagen antibodies. The middle panels show the AVC region and the right panels a high magnification of a subset of cells within the AVC (e) A section of the whole embryo is shown in the left panel; the middle and right panels show the heart and the cells injected in the AVC. The heart sections were stained with anti-periostin antibody. Results are representative of at least 9 experiments. SAC, IAC: superior and inferior AV cushions, PLV primary left ventricle, A atrium. The scale bar indicates 50 μm.

Thus, HPVCs respond in culture to similar cues and use equivalent signaling pathways to undergo EMT as mouse endocardial cells *in vivo*.

Valvular interstitial cells (VIC) give rise to tenocytes and osteo/chondrogenic cells^31,43^. We thus tested the tendinous/chondrogenic potential of HPVCs. We applied for 2 weeks a chondrogenic medium^31^ to HPVCs aggregated in pellets, and found turned-on expression of *SCLERAXIS* and *COLLAGEN1* genes (Fig. 2c) as well as SOX9 and CALCITONIN proteins, suggesting a broad valve repertoire of differentiation of HPVCs (Fig. 2d)

### Human prevalvular cells undergo EMT in *ex vivo* models

To further test the differentiation properties of HPVCs, we used collagen hydrogels to induce EMT. Cells were aggregated in hanging drops for 18 h generating spheroid bodies cultured on collagen gels. SSEA1^-^/MESP1^-^ cells remained at the surface of the gel and proliferated. In contrast, HPVCs invaded the gel and acquired an elongated mesenchymal morphology (Fig. 3a). To confirm that the EMT process led to the generation of valvular interstitial cells, HPVCs were cultured for 18 h within hanging drops and then transferred onto collagen hydrogels. 72-96 h later, gels were fixed and cells immunostained to test expression of post-EMT markers (i.e., smooth muscle actin (SMA), FILAMIN A, PERIOSTIN, and VERSICAN and AGGRECAN antibodies). Figure 3b shows that human cells expressed FILAMIN A, PERIOSTIN, VERSICAN, AGGRECAN and smooth muscle actin (SMA).

Next, we utilized three-dimensional collagen substrate or heart explants^30^ to test the behavior of HPVCs *ex vivo*. AVC (and OFT, data not shown) regions of E9.5 mouse embryos were dissected and placed on a collagen hydrogel (Figure 3c top inset). HPVCs were added to AVC explants for two days, fixed, and stained with anti-SOX9, -FILAMIN A or anti– PERIOSTIN antibody. Figure 3c shows that human cells identified by an anti-human LAMIN A/C antibody and in contact with endocardial cells of the mouse embryonic explant underwent EMT and expressed SOX9, FILAMIN A and PERIOSTIN. One of these cells could be marked by anti-human CD31+ antibody (data not shown)

Thus, HPVCs are capable of recapitulating the EMT process in response to myocardial factors, likely BMP2 in an *ex-vivo* setting.

### Human prevalvular cells undergo EMT *in vivo* in mouse embryos

To investigate the potential of differentiation of HPVCs *in vivo*, we tested whether these cells could further differentiate into valvular fibroblasts when placed in the embryonic environment at a stage of development prior to endoMT. To track HPVCs that have undergone Endo-MT *in vivo* in mouse embryos, a HUES cell line expressing GFP under the transcriptional control of the *Sox9* promoter was engineered. Sox9 was chosen as it is enriched in valves when the cushion endocardial cells were just at the beginning of the EMT process. HPVCs derived from the Sox9 GFP HUES clone were injected into the AVC wall with injections mapped by fluorescent dye (Fig. 4a)^29^. Four to six hours after cell injection, embryonic hearts were dissected and cultured at the liquid-air interface in a matrigel coated insert for 48 h in DMEM added with 50% FCS. Hearts cultured under these conditions maintained their shape and beating activity (data not shown) for 72 h (Fig. 4b). Examination of sections of paraffin-embedded cell-injected hearts revealed activation of the *Sox9* reporter in transplanted ESCs-derived HPVCs. Specificity of the enhancer/promoter was validated by a counterstain of GFP^+^ cells using an anti-SOX9 antibody. HPVCs stained by the anti-human LAMIN A/C antibody remained in the AVC region (Fig.4b).

Alternatively, HPVCs derived from the *Sox9*GFP HUES clone were injected into the AVC of E9.5 embryos and embryos cultured in their yolk sac for 24-48 h. Hearts were allowed to develop in culture (Fig. 4c). The HPVCs injected in the chamber or ventricular wall or in an extra cardiac environment (brain) survived poorly and did not express GFP, SOX9, PERIOSTIN, FILAMIN or COLLAGEN (Fig. 4c, data not shown). Immunostaining of *Sox9*GFP cells revealed that cells expressed PERIOSTIN, FILAMIN A AND COLLAGEN I (Fig. 4d). At high magnification, *Sox9*GFP cells were surrounded by extracellular matrix proteins PERIOSTIN and COLLAGEN. Filamin A (FLNA), which functions as a scaffolding protein and couples cell cytoskeleton to extracellular matrix, was also located at the periphery of the cells (Fig. 4d, right panel). *Sox9*GFP HUES cell derived HPVCs injected in the E9.5 AVC were also found in the AVC both in the superior and inferior atrio-ventricular cushions and expressed periostin following 36 h culture of whole embryos (Fig. 4e).

### Patient specific iPS cells recapitulate features of mitral valve prolapse

To test the pathological relevance of HPVCs and derived valvular cells, we used a patient specific iPS cell model. iPS cells were derived from valvular interstitial cells (VICs) isolated from the explanted mitral valve of a patient harboring a mutation in *Dachsous* gene (*DCHS1*) a likely cause for mitral valve prolapse^6^. In contrast to primary VICs culture, limited by the number of passages, iPS cells provide a mean to recapitulate the stepwise determination of distinct endocardial, valve endothelial, and valve interstitial cell phenotypes. iPS cells allow to repeat separate differentiation experiments both with VECs and VICs. iPs cells expressed OCT4, SOX2 and NANOG (Fig.5a). DNA sequencing confirmed that the patient specific *DCHS1* mutation c-6988C>T was conserved in iPS cells (Fig5b) and digital PCR of copy number variants of main recurrent abnormalities loci found in pluripotent stem cells (technology stemgenomics http://www.stemgenomics.com/) revealed that genomic integrity was conserved (data not shown). Using directed differentiation protocols, cells were capable to differentiate into the three germ layers following treatment with specific morphogens as shown by expression of *SOX17*, and *FOXA2* (endoderm) *BRACHYURY (T), MIXL1*, (mesoderm) and *NESTIN* and *PAX6* (ectoderm) (Fig 5c).

**Figure 5:**
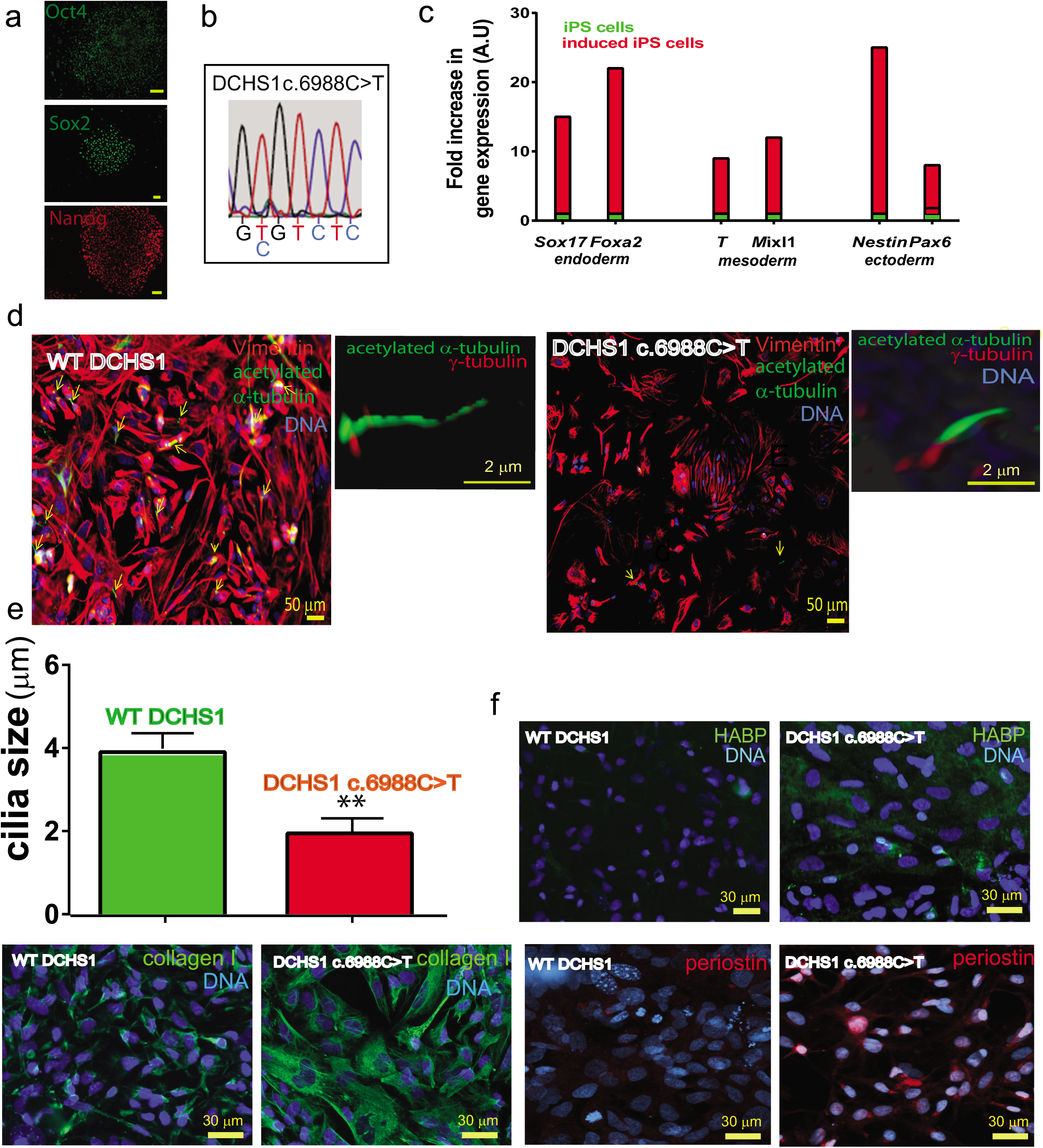
DCHS1 c.6988C>T iPs cells recapitulate the cell pathophenotype of the patient valvular cells. (a) iPS cells derived from mitral valvular interstitial cells express OCT4, SOX2 and NANOG. (b) The patient specific DCHS1 c.6988C>T mutation was conserved following iPS cell derivation. (c) DCHS1 c.6988C>T carrying iPS cells were differentiated toward the endoderm, mesoderm and ectoderm. Real time PCR of *Sox17*,*Foxa2* (endoderm), *Brachyury (T) and Mixl1* (mesoderm) and *Pax6* and *Nestin* (ectoderm) confirmed multilineage propensity. (d) wild type (WT DCHS1) or mutated (DCHS1 c.6988C>T) HPVCs underwent EMT under BMP2 stimulation and were immunostained with anti-vimentin and α-acetylated tubulin to visualize cilia. Insets show 3D-image of cilia co-immunostained with both anti-α-acetylated tubulin and anti-γ-tubulin antibody. (e) Quantification of cilia size: Cilia were stained by anti-α-acetylated tubulin and anti-γ-tubulin. Length of the cilia was measured using Image J in 240 cells from 3 separate experiments. (f) Hyaluronan, collagen I and periostin were visualized by anti-HABP antibody, anti-collagen I and anti-periostin antibodies, respectively in cultures of VIC derived from both wild type (WT DCHS1) or mutated (DCHS1 c.6988C>T) HPVCs. To quantify hyaluronan and collagenI, 3 fields in 3 separate experiments were scored using the image thresholding mode of Image J.

Dachsous is involved in planar cell polarity signaling and in turn cilia formation^44^ and is required for lymphatic valve formation^45^. Valve endothelial cells and their human counterpart iPS cell-derived HPVCs expressing endocardial genes TBX2, *TBX20, PITX2, GATA5, SMAD6* and *MSX1* (Supplementary Fig. 4), expressed few cilia (data not shown) as recently reported in mouse^46^ potentially because the high shear stress in the region of cardiac cushions as proposed in zebrafish^47^. EMT was thus induced by BMP2 treatment of HPVCs and cilia were scored in vimentin+ valvular interstitial cells (Fig. 5d). 450 HPVCs stained with anti-acetylated α-tubulin revealed that 56 % of wild type (wt) cells while only 12% of DACHS1 c.5988 C>T cells featured cilia. The identity of cilia was confirmed by counterstaining with an anti–γ-tubulin antibody (insets Figure 5e). Furthermore, cilia of mutated cells were twice as short as the ones of wild type cells (Fig.5f).

Mitral valve prolapse as a consequence of *DCHS1* c.6988C>T mutation leads to a myxomatous degeneration and an increase in proteoglycan and collagen I expression compared to wild type healthy valve^6^. Wild-type and *DCHS1* c.6988C>T HPVCs derived interstitial cells were thus stained with the hyaluronan-binding protein to visualize hyaluronan and an anti-collagen I antibody. Figure 5f revealed that mutated cells secrete three times as much collagen I (22.7±3 % of cell field, n=5 experiments, 540 scored cells) than wild type cells (7.4±1.2 %) and six times as much hyaluronan (36± 6%, of cell field n=5 experiments, 645 scored cells) than wild-type cells (5.7±1.5 %). They also featured 7 times as much periostin (28 ±4%, n=5 experiments, 530 scored cells) than wild type cells (4±0.8%).

### Sonic hedgehog (SHH) a targeted signaling pathway to prevent *DCHS1* c.6988C>T patient specific valvular cells from over-secreting ECM proteins

SHH is a signaling pathway of primary cilia. SHH regulates expression of hyaluronan^48^, and play key roles in ECM remodeling in development^49^. SHH receptor Patched1 is located on or at basal body of cilia and relocates to the entire cell membrane when bound by its ligand where it gets activated even without ligand^50^. We reasoned that DCHS1 c.6988C>T valvular cells with short or no cilia may feature, following a lack of Patched1 on cilia, a constitutive activation of SHH pathway. We first looked at the location of Patched1 in wt and DCHS1 c.6988C>T cells. Figure 6a shows that Patched1 was clustered and aggregated at the cilia basal body in wt cells while it was spread as clusters all over the cell membrane in non-ciliated DCHS1 c.6988C>T.

**Figure 6:**
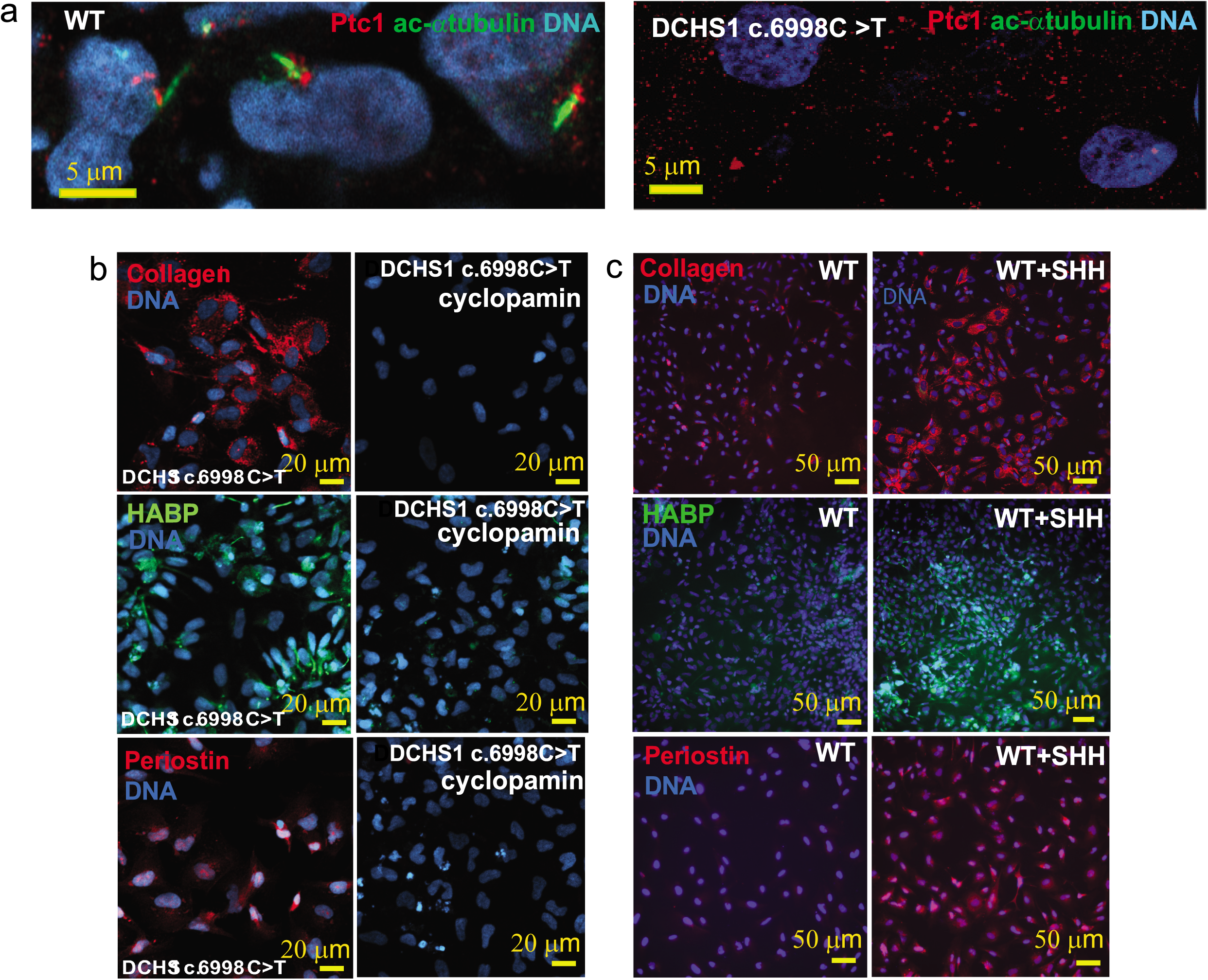
SHH overactivation in mutated (DCHS1 c.6988C>T) cells. (a) SHH receptor patched I (Ptc1) and the cilia (anti-acetylated α-tubulin) were co-stained on both ciliated wild type (WT) or unciliated DCHS1 c.6988C>T cells. The confocal images are representative of 3 separate experiments and 180 scored cells in each group. Hyaluronan, collagen I and periostin were visualized by anti-HABP antibody, anti-collagen I and anti-periostin antibodies, respectively in cultures of VIC derived from both mutated (DCHS1 c.6988C>T) or wild type (WT DCHS1) HPVCs treated with cyclopamin (b) or SHH (c), respectively. To quantify collagen I, hyaluronan and periostin, 3 fields in 5 separate experiments were scored using the image thresholding mode of Image J.

We thus treated DCHS1 c.6988C>T cells with cyclopamine, a hedgehog pathway inhibitor, together with BMP2 at the onset of EMT. The drug prevented in a significant manner (p≤0.01) secretion of excess collagen I (from 25±3 down to 2±0.3% cell field, n=3, 520 scored cells) hyaluronan (from 30±2.1 down to 1.65±0.5% cell field, n=3, 480 scored cells), or periostin (from 20±1.7 down to 1.5±0.2% cell field, n=3, 515 scored cells) in DCHS1 c.6988C>T cells (Fig. 6b). We next tested whether SHH added to wt cells could mimic DCHS1 c.6988C>T phenotype. Addition of 100 ng/ml SHH for 48 h on wt cells treated with BMP2 increased in a significant manner (p≤0.01) expression of collagen I (from 6.8±1 up to 16.7±2.1% of cell field, n=3, 340 scored cells), hyaluronan (from 10±1.2 up to 17.1±1.9% of cell field, n=3, 380 scored cells) and periostin (from 4± 0.7 up to 31.2± 2.7% of cell field, n=3, 490 scored cells) (Fig. 6c). Interestingly in contrast to wild type cells (Fig.7), single cell sequencing of DCHS1 c.6988C>T post-EMT cells revealed that cells were dramatically heterogeneous as illustrated by the t-SNE graph (Fig.7a) and the heatmap of cell cluster (Fig 7c). A principal component analysis of WT *vs* DCHS1 c.6988C>T single valvular interstitial cell transcriptome revealed a distinct pattern of gene expression (Fig.7b). Differences between WT and *DCHS1* c.6988C>T cells could not be attributed to a differential cell cycle as most cells of the two populations were in the G1 stage (inset Fig.7b).

**Figure 7:**
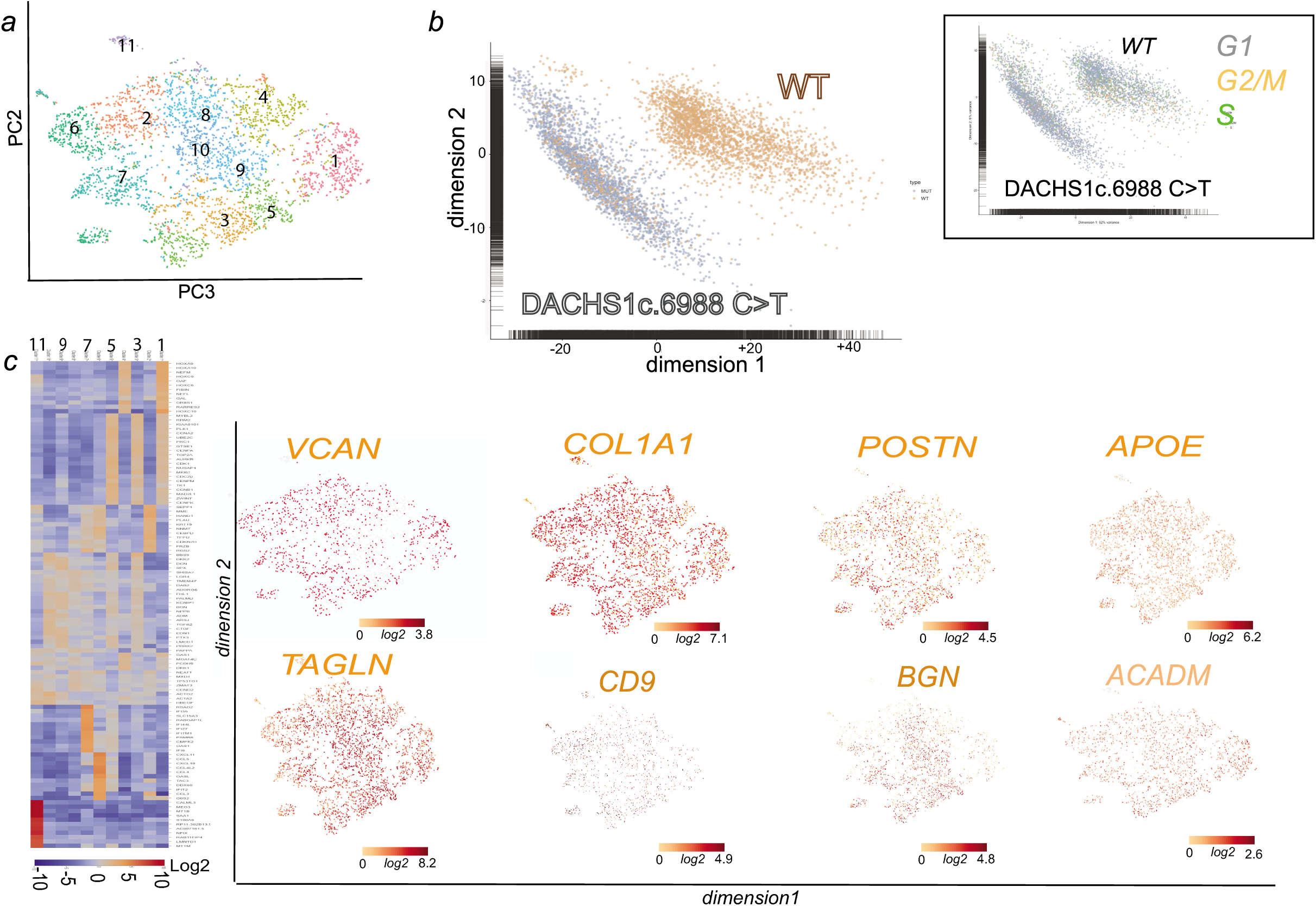
Single cell-sequencing of DCHS1 c.6988C>T post-EMT valvular insterstitial cells. (a) t-distributed stochastic neighbor embedding (t-SNE) 2D cell map 10X genomics (n=4316 cells) (upper panel).). (b) PCA of wt versus DCHS1 c.6988C>T post-EMT (BMP2 treated) HPVC cells (c) Graph-based Log2 fold changes in gene expression of cell clusters compared to all other cells (left panel) and Highlight of cell populations expressing genes marking fibrosa or spongiosa valve layers.

Cells could not acquire a phenotypic identity. All expressed high level of *TAGLN*, collagen I (*COL1A1*), Versican (*VCAN*) and periostin (Fig. 7). Some genes enriched in a specific valve layer such as TDGF1 (fibrosa) were not expressed at all in *DCHS1* mutated cells or like *CD9, APOE, BGN* (fibrosa) or *ACADM* (spongiosa) spread all over cell clusters (Fig. 7c). A small cluster (cluster 6) of cells were endothelial cells highly expressing *PECAM1, CDH5, ENG, SOX17, SOX18, DLL4, CD34 and KDR* (Supplementary table 4). Interestingly GLIs 2,3 were also overexpressed (9.3±1.1 fold in 3 real-time PCR experiments and in single cell datasets DCHS1 c.6988C>T vs wt cells (see Supplementary table 3 and 4 and supplementary Fig 5). Furthermore *BRD2* and *PRTM1*, two positive regulators of SHH ^51^ were highly upregulated and more broadly expressed in DCHS1 c.6988C>T vs wt valvular cells ‘supplementary Fig.5).

## DISCUSSION

The present study documents the propensity of pluripotent stem cell-derived MESP1^+^ cardiovascular progenitors to give rise to *bona fide* HPVCs which in turn provide a reliable source for functional signal-responsive valvular cytotypes, namely valvular interstitial cells and tendinous/chondrogenic cells. Establishment of pluripotent stem cell derived HPVCs offers a powerful prototype platform of human early valvulogenesis addressing a major gap in this field. The ensuing application of patient specific iPS cell-derived valvular cells to model replay valvulopathy exemplifies a genuine model of patient-specific nonsyndromic mitral valve prolapse underscoring the biological validity and clinical utility of this proof-of-concept study.

### Novel model to study human valvulogenesis

To map the gene expression signature for the HUES-cell derived valvular progenitor cell population (HPVC), we performed gene expression profiling on one of the region of the heart where the valves form. We used E9.0 embryos to determine the transcriptomic identity of AVC prior to EMT, which occurs between E9.5 and E11. We show here that HPVCs derived from pluripotent stem cells accurately reflect the *in vivo* E9.0 AVC profile. The gene profile of AVC was compared to the adjacent primary chamber myocardium and endocardium, thus restricting the cell gene signature to the endocardium of the AVC. While the primary ventricle may feature to some extent different gene profile than the AVC myocardium, our primary ventricle gene array did not show expression of *CKM*, a gene not expressed in AVC myocardium in contrast to mature chamber myocardium^52^ pointing to a still immature myocardium. Gene expression arrays of this cardiac-restricted region (i.e., using Tbx3 GFP sorted cells from E10.5 mouse embryos) so far available in the literature were done at a later stage of development, thus during EMT, and included both endocardium and myocardium^53^. An attempt to carry out a transcriptome of E9.5 AVC was reported previously^38^. However, a careful analysis of these data revealed that the AVC transcriptome was contaminated by high expression of chamber specific genes such as *TnT*, or *Gja1* (see GEO data set GDS3663) and key AVC specific genes such as *Tbx2* were missing (see GEO dataset GDS3663 and Figure 1D)). This could explain why this previously reported gene signature did not cluster tightly with those of our dissected AVC and HPVCs (Figure 1E). One study using a SAGE protocol reported a gene expression profile of mouse cardiac AVC at a slightly later stage (E9.5) of development^54^. The comparison of our arrays data with that of the Tbx3-GFP-sorted cells^53^ and the SAGE data^54^ further revealed the expression of common and major genes in AVC *vs.* cardiac chambers. These include *Tbx2, Tbx3*, *Tbx20*, *Msxs, id2*. Interestingly, four of these genes (*TBX3, TBX2, TBX20,* and *MSX*) were highly expressed in the HPVCs (Figure 2). However, our array identified additional endocardial-specific genes, like *GATA5*, or *NFATC*^55^ as well as genes restricted to AVC such as *TGFβ2,* and *HEY1* ^42,56^ also expressed in the HPVCs including at a single cell level (Figure 1). Genes suggestive of the cardiac cushions such as *Smad6*, *Twi*st, endocardial *Msx*^*57*^, a serotonin receptor variant *(Slc6a4)*^58^ were also found enriched in AVC endocardium *vs* chambers. Furthermore we uncovered genes highly enriched in the AVC that were not previously reported to play a role in AVC identity or more specifically in cushion formation were. These include caveolin (*Cav1*), specifically expressed in the endocardium (Figure 1) thrombospondin (*Thsd*2), the Wnt modulator R-Spondin (*Rspo3*), the adenosine receptor (*Adora*) and the neuropeptide galanin (*Gal*)^59^. Expression of most of these genes was also found in the HPVC population (Figure 2). None of the genes found enriched in expression in the chambers (*Nppa, Irx2*) were present in AVC, nor in the HPVCs. Single cell data further confirmed previous data. At a single cell level, *endoglin (ENG)* an endocardial specific gene, *VE-Cadherin (CDH5), KDR* and *PECAM1, THSD1*were found highly enriched et different levels either in *TWIST1* negative or positive cells. GAL, ADORA1, and CAV1 were all expressed at a single cell level.

The gene profiles of HPVCs as well as AVC were quite different from the one of either the inferior or superior outflow tract. None of the main enriched genes in both AVC and HPVCs (*Cav1, Thsd1, Spo3, Vsnl1, Gal, Igfbp7, Shisa2, Tbx2, hey1, wnt2, Pitx2…*) were expressed in OFT regions^60^ thus pointing to a specific AVC print of HPVCs.

These findings collectively point out to the pre-valvular endocardial identity of the human cell population and validate the protocol used to derive such a cell population. Overall, the transcriptome of HPVCs clustered with the one of E9.0 mouse AVC and not with mesenchymal cells (Figure 1), further emphasizing the identity of the HPVC. Together, these novel data reveal the unique potential of our HPVC model to discover and further investigate regulation of human valvulogenesis.

### Developmental origin of valve precursor cells

Our rationale to direct the fate of MESP1^+^ cardiovascular progenitor towards the myocardium and endocardium, and further to the endocardial cushions^61,^ and a valvular fibroblastic as well as tendinous/chondrogenic phenotype, was that these progenitors should acquire an endothelial identity and an endocardial cell fate within a cell population at the expense of myocardial outcome in line with valvular prioritization. We have found that FGF8, known to favor endocardium *vs* myocardium^34^ as well as in the formation and EMT of cardiac cushion^63–65^, was potent in recapitulating the same cell determination process from pluripotent stem cells-derived MESP1^+^ cells^25^. We thus combined the action of VEGF, an endothelial inducer, with FGF8 driving mesodermal cell fate toward endocardial cells at the expense of myocardial lineage^34^. Stimulation by VEGF and FGF8 were combined with the influence of extracellular fibronectin matrix and MEF, which secrete FGF2 and TGFβ 2^66^, known to synergize with VEGF to determine the endocardium^67^. Together, this inductive signaling induced the HPVC to express endothelial (i.e *CD31, VE-CADHERIN, ENDOGLIN*) and endocardial lineage genes as found in the AVC (i.e. *GATA5*^+^, *TBX2*^*+*^, *GALANIN*^*+*^, *RSPO*^*+*^, *SMAD6*^*+*^, *MSX1*^*+*^ cells). This suggests that HUES cells can not only recapitulate the mesodermal cardiogenic pathway ^25^ but can also be directed toward the fate of endocardial pre-valvular AVC cushion cells (Fig. 6). These findings correlate with embryonic development as MESP1^+^ cells are known to give rise to endocardial cells and cardiac cushions in the mouse embryo^19,62^.

### HPVCs as a model for endocardial cushion cells

We further show that human valve progenitors are capable of EMT under different experimental conditions. First, when cultured on collagen gels, HPVCs acquire a mesenchymal phenotype and express valvular fibroblast markers such as FILAMIN, PERIOSTIN, VERSICAN and AGGRECAN. Second, when cultured *ex vivo* or grafted *in vivo* with or close to cardiac cushions of the AVC in mouse embryos, respectively, only HPVCs, but not SSEA1- or SSEA1+ MESP1+ cells, become valvular interstitial cells expressing FILAMIN A, PERIOSTIN, SOX9 and COLLAGEN I, indicating that the HPVCs are unique and selective in their response to cues derived from the surrounding mouse AVC tissue.

The EMT process is likely triggered *ex-* and *in-vivo* through the secretion of BMPs by the myocardium, as indicated in *in vitro* mouse AVC explant cultures^68^ and in genetically altered *in vivo* mouse systems^38, 68^. Such a phenomenon could be recapitulated *in vitro* using BMP2 and notch signaling pathways activation with HUES cells-derived HPVCs. Indeed, BMP2 also induces expression of post EMT genes such as *SLUG, SMAD6, PERIOSTIN,* or *SOX9*. Expression of *SLUG*, a direct notch target *in vitro* and *in vivo*^42,69^, and *PERIOSTIN* were abrogated by the Notch inhibitor, DAPT. *PERIOSTIN*, *SLUG* and *MSX1* expression was further induced by ectopic expression of the intracellular Notch domain (Supplementary Fig. 2). These findings thus confirm that maturation of stem cell derived HPVCs recapitulates *in vitro,* at least partially, the *in vivo* embryonic signaling mechanisms mediating EMT and cell invasion^70^, including the interplay between the Bmp2/Notch signaling pathways^42^.

### HPVCs as a model of valvulopathies

Primary ciliary dyskinesia (PCD) without sinus invertus as well as other ciliopathies have been clinically associated with myxomatous mitral valve and other valve diseases^71,46^ Interestingly, patients with autosomal dominant polycystic kidney disease (ADPKD), a disease in which cells feature a over-activation of SHH pathway^72^ have an increased occurrence of mitral valve prolapse^73^. Dachsous is required for lymphatic valve development^45^. Dachsous mutated HPVCs recapitulated several features of mitral valve prolapse (Fig. 5). This included cilia defects responsible for disorganization of valve interstitial cells within the leaflets and excess secretion of hyaluronan, collagen I and periostin, a hallmark of mitral valve prolapse.

HPVC single cell-data uncovered that *DCHS1* c.6988C>T valve interstitial cells could not acquire normal identity and express at high levels tissue-specific *COL1A1* or *PSTN*. This lack of cell identity could explain that the cells could not properly distribute within valve layers. This also suggests that ECM may regulate cell fate of valve interstitial cells in the course of leaflet patterning. Interestingly, cilia signals through sonichedghog that targets through the transcription factor Gli3 the hyaluronic acid synthetase (*Has*) gene^74^ and other components of the ECM^49^. In the absence of sonic hedgehog (SHH), the GLI proteins GLI2 and GLI3 are phosphorylated by PKA, CKI and GSK3β. This leads to their proteolytic cleavage to generate repressor forms (GLI2R and GLI3R, respectively)^75^. A deregulation of this signaling pathway would relieve inhibition on *Has* and up-regulate hyaluronic acid. Besides the role of cilia as a sensor of the extracellular environment (i.e. EC Matrix), the direct regulation of ECM organization may explain that mutation in *Dachsous* and absence of cilia deregulate synthesis of ECM proteins through SHH pathway. Indeed, SHH receptors patched1 were found clustered at the basal body of cilia in wild type cells while were spread on the cell membrane in *DCHS1* c.6988C>T cells lacking cilia (Figure 6). Furthermore GLIs 2 and 3 as well BRD2 and PRTM1, positive regulators of SHH targeted genes^51^ were upregulated in *DCHS1* c.6988C>T cells valvular cells vs wild type cells pointing to an activation of SHH pathway. Interestingly, a lack of cilia favors EMT in mouse Tg737/ift88 mutant^76^ in agreement with our observation that *DCHS1* c.6988C>T undergo EMT as wt cells. However, *DCHS1* c.6988C>T mutant featured an over-activation of SHH pathway in contrast to Tg737/ift88 mutant heart which lacked SHH signaling at least in second heart field progenitors^76^.

Our data using a SHH inhibitor (Figure 6) confirmed that SHH signaling is constitutively activated in *DCHS1* c.6988C>T cells. Regulation of the pathway by a pharmacological SHH pathway inhibitor rescued the ECM cell phenotype. Thus, our data bring a potential pharmacological target to potentially prevent MVP.

This proof of concept application underscores that patient-derived iPS cells can be used as faithful models in a dish of human valve diseases.

### Future considerations

A limitation of our study was the use of SSEA1^+^ MESP1+sorted cells. Such sorting may have excluded the endocardial cells that are already determined in the primitive streak^13,15,77^ and segregated from the myocardial lineages in the pre-cardiac mesoderm prior to the segregation of the first and second myocardial lineages^14,67^, as reported in chicken, quail or mouse embryos. Whether, at least some endocardial cells are already determined in the pre-cardiac human mesoderm remains to be further investigated.

Our *in vitro*, *ex vivo* and *in vivo* assays facilitate further studies examining the events that give rise to the human endocardium, valvular interstitial cells, and tendinous/chondrogenic cells, as well as to delineate the pathways triggering EMT. This human model of valvulogenesis could be extended to patient-specific iPS cells and will help in understanding the molecular mechanisms underlying valvulopathies originating from defects in cell fate decisions and/or EMT.

## Acknowledgments

We are grateful to the UCSD Neuroscience Microscopy Shared Facility for access to the Spinning disk microscope, to Jeff Smith and Dr Joan Heller Brown (UCSD, dept of Pharmacology, UCSD) for kindly allowing us to use their microinjection set up, and to Daniel Stockolm (Genethon, Evry) for access to the cell imaging facility. We are also Thankful to Dr Valérie Balme and colleagues at HALIODX (Marseille) for helpful discussions and skillful experiments of single cell sequencing.

